# p53 is not necessary for DUX4 pathology

**DOI:** 10.1101/118315

**Authors:** Darko Bosnakovski, Erik A. Toso, Olivia O. Recht, Anja Cucak, Abhinav K Jain, Michelle C. Barton, Michael Kyba

**Affiliations:** Lillehei Heart Institute, University of Minnesota, 312 Church St. SE, Minneapolis, MN 55455; Department of Pediatrics, University of Minnesota, 312 Church St. SE, Minneapolis, MN 55455; University Goce Delcčev - Sštip, Faculty of Medical Sciences, Krste Misirkov b.b., 2000 Sštip, R. Macedonia; Department of Epigenetics & Molecular Carcinogenesis, The University of Texas MD Anderson Cancer Center, Houston, TX 77030

**Keywords:** facioscapulohumeral muscular dystrophy, DUX4, p53

## Abstract

FSHD is a genetically dominant myopathy caused by mutations that cause expression of the normally silent *DUX4* gene. This transcription factor has been shown to interfere with myogenesis when misexpressed at very low levels in myoblasts, and to cause cell death when overexpressed at high levels. A previous report using adeno-associated virus to deliver high levels of DUX4 to mouse skeletal muscle demonstrated severe pathology that was suppressed on a p53 knockout background, implying that DUX4 acted through the p53 pathway. Here, we investigate the p53-dependence of DUX4 using both in vitro cellular and the transgenic iDUX4[2.7] mouse models. We find that inhibiting p53 has no effect on the cytoxicity of DUX4 in vitro. When crossed onto the p53 null background, we find no suppression of the male-specific lethality or skin phenotypes of the DUX4 transgene, and find that primary myoblasts from this mouse are still killed by DUX4 expression. These data challenge the notion that the p53 pathway is central to the pathogenicity of DUX4.

**Summary Statement:** DUX4 is thought to mediate cytopathology through p53. Here, DUX4 is shown to kill primary myoblasts and promote pathological phenotypes in the iDUX4[2.7] mouse model on the p53-null background, calling into question this notion.

## Introduction

Facioscapulohumeral muscular dystrophy (FSHD) affects over 25,000 people in the U.S. alone, making it one of the most prevalent genetic diseases. The genetic mutation underlying FSHD is usually a reduction in the copy number of a macrosatellite repeat on chromosome 4 referred to as D4Z4 (van Deutekom et al. 1993;Wijmenga et al. 1992). This repeat is GC-rich, highly methylated, and normally subjected to repeat-induced silencing, which is disrupted in an allele-specific manner by contractions to 10 or fewer copies (van Overveld et al. 2003) or on all D4Z4 repeats by mutation in the chromatin protein *SMCHD1* (de Greef et al. 2009;Hartweck et al. 2013;Lemmers et al. 2012). When silencing at D4Z4 breaks down, an RNA transcript encoding the DUX4 protein (Gabriels et al. 1999) is expressed. The presence of a poly(A) signal downstream of the D4Z4 repeats on chr4 (Dixit et al. 2007) leads to DUX4 expression and explains why disease is associated only with contractions of the D4Z4 repeats on chr4, and only on specific 4qter alleles, so called permissive alleles, which harbor the poly(A) signal (Lemmers et al. 2010;Lemmers et al. 2004). The DUX4 protein has been observed by immunostaining of cultured FSHD myoblasts which show infrequent, possibly episodic, expression (Jones et al. 2012;Snider et al. 2010) and differentiated myotubes which show more prevalent expression (Block et al. 2013;Rickard, Petek and Miller 2015).

Mechanisms of DUX4-mediated pathology are currently being investigated actively. At high levels of expression, DUX4 causes toxicity that leads to cell death of myoblasts (Bosnakovski et al. 2008;Kowaljow et al. 2007), while at low levels of expression, it impairs myogenic differentiation (Dandapat et al. 2014) and sensitizes cells to oxidative stress (Bosnakovski et al. 2008). A mouse model allowing doxycycline-regulated DUX4 expression has recapitulated these phenotypes in primary myoblasts (Dandapat et al. 2014) while also having several nonmuscle related phenotypes due to low basal levels of DUX4 expression in the absence of doxycycline (Dandapat et al. 2016). Other animal model work includes a mouse carrying human D4Z4 repeats (Krom et al. 2013) which showed some evidence of sporadic DUX4 expression, but no myopathy; and adeno-associated viral (AAV) vector-mediated delivery of DUX4 to skeletal muscle, which showed profound myopathy (Wallace et al. 2011). This latter work implicated the p53 pathway in DUX4 pathology, as the p53 knockout background suppressed AAV-DUX4 toxicity. The linkage to p53 is compelling, as this pathway has the potential to push cells into apoptosis, however precisely how DUX4 would activate p53 is not clear. The immediate targets of the DUX4 transcription factor include genes with regulatory elements containing the sequence TAATCTAATCA (Geng et al. 2012;Zhang et al. 2015), or variants thereof. ChIP-seq has identified many genomic targets (Choi et al. 2016;Geng et al. 2012) and at the majority of these, DUX4 displaces nucleosomes, recruits p300 and/or CBP through its C-terminus which promotes acetylation of histone H3 and activation of transcription (Choi et al. 2016).

Here, we directly test the p53-dependence of the pathogenic phenotypes of DUX4 expression, by inhibiting p53 in cell lines while DUX4 is expressed, and by crossing the iDUX4[2.7] transgene onto the p53 knockout background. These experiments reveal that p53 is not necessary for DUX4-mediated pathology, neither in myoblasts, nor in other tissues.

## Results

Previously, overexpression of Pax3 or Pax7 was been shown to reduce the toxicity of DUX4 to C2C12 mouse myoblasts (Bosnakovski et al. 2008). To see to what extent inhibiting p53 could similarly reduce DUX4 toxicity, we transduced iC2C12-DUX4 cells, which are immortalized mouse myoblasts engineered for doxycycline-inducible DUX4 expression, with retroviral vectors expressing several constructs known to interfere with p53: a dominant negative p53 (R175H mutation), MDM2 (Momand et al. 1992), and TRIM24 (Allton et al. 2009). Overexpressing cell lines were established and cells were exposed to two different doses of doxycycline, both of which resulted in significant cell killing in the absence of overexpression (empty vector control). Notably, although the positive controls, *Pax3* and *Pax7,* protected substantially at 24 hours, cells transduced with the three p53 inhibitors did not show a reduced death rate compared to empty vector (Fig. 1). This indicates that in this system, p53 is not a relevant player in the cell death phenotype.

**Figure 1.**
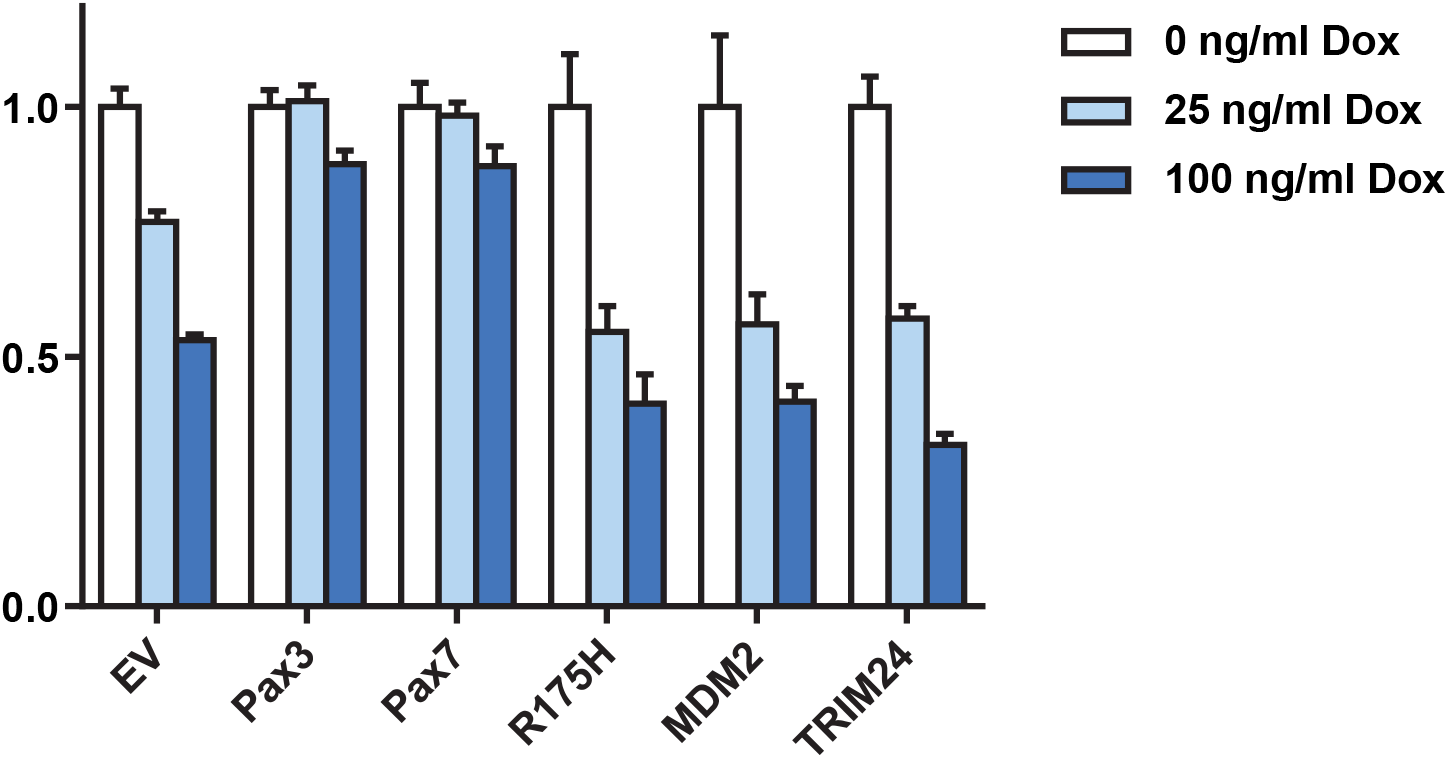
Inhibiting the p53 pathway has no effect on viability in vitro. Viability of iC2C12-DUX4 cell lines which express DUX4 in response to different doses of doxycycline and other factors constitutively from the MSCV LTR. Viability in the presence of doxycycline is normalized to that in its absence.

We have previously generated a mouse model with a doxycycline-inducible 2.7 kb DUX4 transgene (Dandapat et al. 2014;Dandapat et al. 2016), referred to as iDUX4[2.7]. One of the interesting features of this mouse is that the very low basal level of expression from the Tet-on promoter in the absence of doxycycline leads to various non-muscle phenotypes, especially in males, where 80% die as embryos, with the remaining 20% being severely runted and all dying before 6 weeks of age. Females are less severely affected and can thus propagate the strain - because the transgene is X-linked, X-inactivation diminishes the phenotype in females. We reasoned that if p53 were necessary for the pathological effects of DUX4 on embryonic cell types, then on a p53 knockout background, males ought to be born at normal ratios, and ought to be relatively healthy compared to siblings with a functional copy of p53. We therefore crossed the iDUX4[2.7] transgene onto the p53 knockout background. Female carriers were obtained that were homozygous for the p53 knockout and these were bred to male p53 heterozygotes. We genotyped 65 progeny from this backcross and obtained no DUX4+ males, neither on the p53 knockout nor on the heterozygous background (Fig. 2). Thus, absence of p53 has no effect on the pathological effects of DUX4 on development in the iDUX4[2.7] strain.

**Figure 2.**
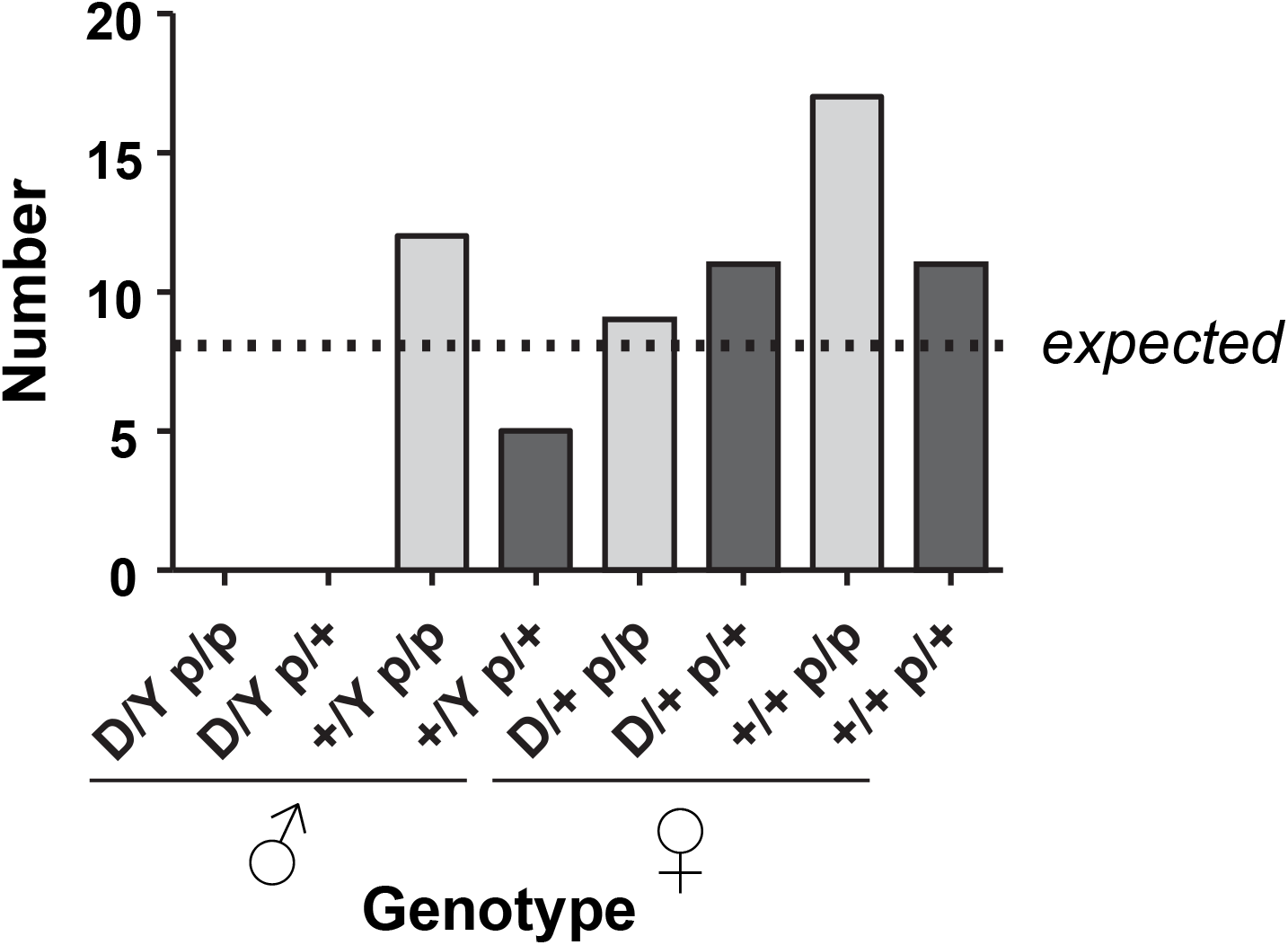
p53 status does not impact survival of iDUX4[2.7] animals to birth. Summary of genotypes observed from a backcross of p53 knockout males to p53 heterozygous females carrying the iDUX4[2.7] transgene. Note that no males carriers were observed, neither in the heterozygous nor homozygous p53 null state. Testing the hypothesis that p53 affects survival, using the 4 classes of male progeny, p=0.74 (Fisher’s exact test), therefore the null hypothesis (p53 does not affect survival of iDUX4[2.7] mice) is assumed. D=iDUX4[2.7] transgene, p=p53 null.

Although no males were produced from the backcross described above, we did eventually obtain live born p53 iDUX4[2.7] knockout males from F1 crosses to Rosa-rtTA p53 double heterozygous males, and these rare animals displayed the severe runting and skin phenotypes (flaky skin, alopecia, and puffy eyelids) typical of the iDUX4[2.7] mouse (Dandapat et al. 2014). Thus p53 is not necessary for the pathological effects of DUX4 on non-muscle tissues.

To determine whether DUX4 would be cytotoxic to muscle cells in the absence of p53, we established primary cell cultures from muscle tissue of these iDUX4[2.7]; p53 KO animals. Primary muscle cells were sorted into myogenic and fibro-adipogenic fractions by flow cytometry for VCAM/Itga7 and PDGFRα, respectively. These sorted primary cultures were then exposed to doxycycline over a series of doses to induce DUX4 expression to different levels. DUX4 expression was clearly cytopathic (Fig. 3A) and caused a dose-dependent loss of cellular viability in myoblasts (Fig. 3B). The same was observed for fibroadipogenic progenitors (Fig. 3B) on the p53 knockout background. Thus, in the absence of p53, DUX4 is still cytotoxic, to both myogenic and fibroadipogenic progenitors.

**Figure 3.**
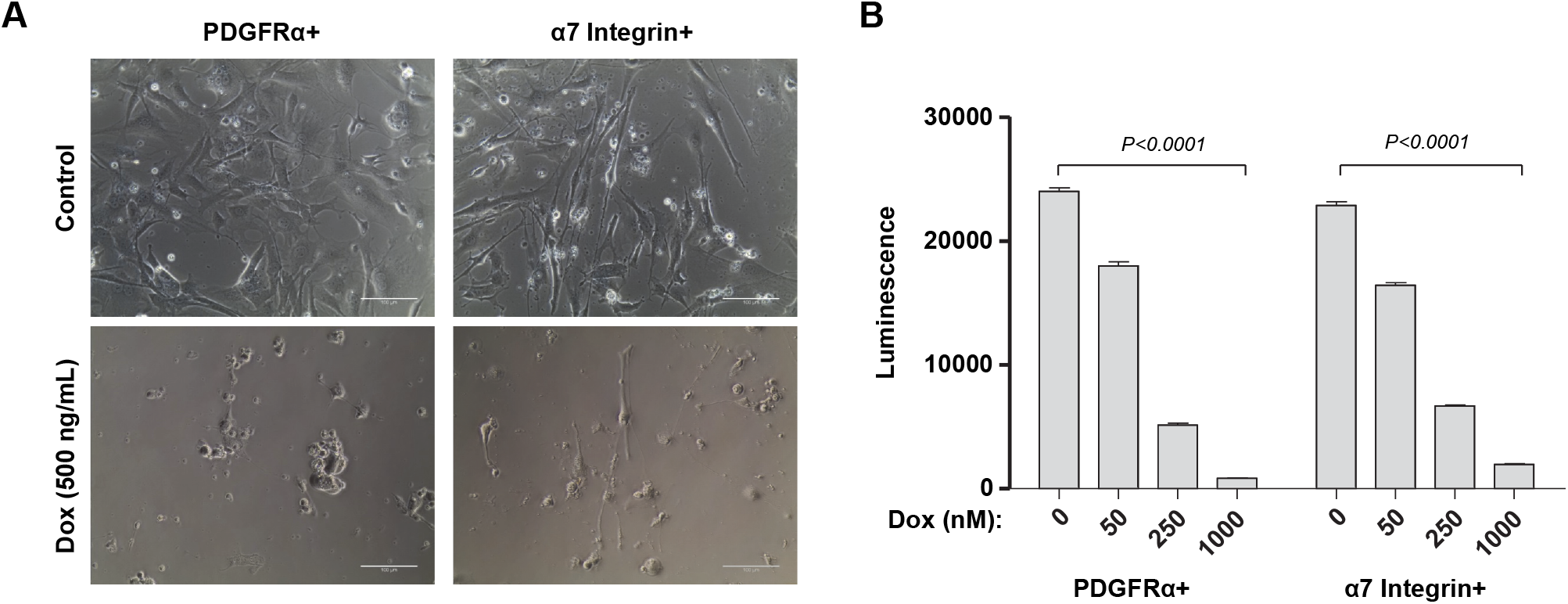
Muscle progenitors are sensitive to DUX4 on the p53 mutant background. (A) Photomicrographs of homozygous fibroadipogenic (PDGFRα+) and myogenic (Integrin α7+) progenitors from male p53-KO; iDUX4[2.7] mice under control expansion conditions (above) or exposed to doxycycline (dox) to induce DUX4 expression (below). Scale bar represents 100 μm. (B) Viability assay for the same cells, exposed to different doses of doxycycline.

## Discussion

Using both the inducible C2C12 cell culture system, where it was originally demonstrated that DUX4 was pathogenic to myogenic progenitors (Bosnakovski et al. 2008), and the iDUX4[2.7] mouse model (Dandapat et al. 2014), we find no evidence for p53 involvement in DUX4 pathogenicity. Interestingly, this applies to phenotypes caused by very low basal levels of DUX4 expression, as the developmental effects and runting are thought to be, as well as overt cytotoxicity in primary or immortalized cells, which is the result of high level DUX4 expression. The previous study implicating p53 in DUX4 toxicity made use of a very different experimental system with vastly higher levels of DUX4 expression: recombinant adeno-associated virus vector-mediated production. The levels of DUX4 expression achieved in a cell infected with multiple copies of DUX4 driven from the CMV promoter are much higher than those in which DUX4 is expressed from its endogenous promoter from a single (terminal) D4Z4 repeat in FSHD. It is possible that at these exceptionally high levels of expression, the p53 pathway does in fact become necessary for some aspect of extreme pathology, however the relevance of such high levels of expression is debatable. The nature of the muscle damage seen in the AAV model is quite distinct from that seen in FSHD, and DUX4 expression is in fact quite difficult to detect in biopsies of FSHD muscle, to the point where the most reliable readout of its expression in biopsies is its target gene fingerprint (Yao et al. 2014). It is also theoretically possible that some aspect of the AAV experimental system combines with the effect of DUX4 to make the pathology more severe than it would otherwise have been, and the p53 effect ascribed to DUX4 was in fact downstream of the cellular response to AAV infection. Nevertheless, the studies presented here clearly demonstrate that p53 is not necessary for cytotoxicity of DUX4.

## Materials and Methods

### Mice

Animal work was conducted under a protocol approved by the University of Minnesota IACUC. The p53 mutant allele used in these studies was obtained from Jackson Labs (strain 008651) and carries a floxed stop codon upstream of a dominant negative (R270H) p53 mutant. In the absence of cre, no functional p53 is produced, thus as used in the current study, it is a null allele. Mice homozygous for this allele developed papillomas and most eventually succumbed to thymic lymphomas, as predicted for a p53 null.

### Cell Culture

The iC2C12-DUX4 cell line was cultured in high glucose Dulbecco’s Modified Eagle Media (DMEM) supplemented with 20% fetal bovine serum (FBS, Atlanta Biologicals), L-glutamine and sodium pyruvate (Gibco), penicillin and streptomycin (P/S, Gibco) at 37 °C in 5% CO2.

Primary mouse myoblasts and muscle fibroblasts were isolated and expanded as previously described (Arpke et al. 2013;Dandapat et al. 2014). Briefly, hind limb muscles were dissected under sterile conditions, minced using razor blade and digest with collagenase type II (Gibco; Grand Island, NY, 17101-015) and dispase (Gibco, 17105-041). Single cell suspensions were plated in T75 dishes in F-10/Ham’s (Hyclone) medium containing 20% FBS (HyClone), 50 ng μL^−1^ human basic fibroblast growth factor (Peprotech), 1% penicillin/streptomycin (Gibco), and 1% Glutamax (Gibco) and cultured at 37°C at 5% O_2_, 5% CO_2_, for 5 days. From the expanded primary cells, myoblasts and fibroblasts were separated by flow cytometry for α7 integrin and PDGFRα integrin expression, respectively. Cells were trypsinized (0.25%, Gibco) and stained with PDGFRα PE (e-Biosciences, clone: APA5) and α7 integrin APC (Ablab, R2F2) antibodies in FACS staining medium (PBS, 1% FBS). Single positive FACS sorted populations were expended in the same medium and conditions used for initial primary culture.

### Expression constructs

We constructed the expression vector pMSCViGFP.GW, a gateway cloning-compatible retroviral expression vector by subcloning the gateway exchange cassette (Invitrogen) into XhoI/BglII-cut pMSCV-ires-GFP. The cDNAs for p53R175H, TRIM24, and MDM2 were amplified by PCR with flanking *attB* sites and subcloned by gateway cloning into MSCViGFP.GW.

### Establishing overexpressing cell lines

Retroviral expression constructs were co-transfected with the packaging constructs pCL-Eco (Naviaux et al. 1996) into NIH-3T3 cells using FuGENE 6 transfection reagent (Roche). Virus-containing supernatants were collected 48 hours later, filtered (0.45 μm) and used directly to transduce iC2C12-DUX4 cells. Constitutively overexpressing cell lines were obtained by FACS sorting GFP+ cells.

### Cell death inhibition assays

Cell viability was analyzed as previously described (Bosnakovski et al. 2014). Briefly, myoblasts and fibroblasts were plated in 96 well plates at 3000 cells per well in proliferation medium and treated with different concentrations of doxycycline for 72 h. Medium was removed from the plate; 100 μL working (diluted 1:3 with PBS) CellTiter-Glo luminescent assay reagent (Promega) was added to each well and luminescence measured on a Cytation3 plate reader (BioTek).

## Acknowledgements

We thank the Dr. Bob and Jean Smith Foundation for their generous support. We thank Micah Gearhart for helpful advice.

## Competing Interests

The authors declare no competing interests.

## Author Contributions

Conceptualization and methodology: MK and DB; writing: MK and DB; reviewing and editing: MK, DB, AKJ, MCB; data acquisition and analysis: DB, EAT, OOR, AC, AKJ.

## Funding

This work was supported by the National Institute of Arthritis and Musculoskeletal and Skin Diseases (R01 AR055685).

